# Deep sequencing of the T cell Receptor reveals common and reproducible CD8^+^ signatures of response to checkpoint immunotherapy

**DOI:** 10.1101/2023.02.11.528080

**Authors:** Robert A. Watson, Chelsea A. Taylor, Orion Tong, Rosalin Cooper, Elsita Jungkurth, Piyush Kumar Sharma, Bethan Storey, Weiyu Ye, Bo Sun, Alba Verge de los Aires, Flavia Matos Santo, Isar Nassiri, James J. Gilchrist, Eleni Ieremia, Mark R. Middleton, Benjamin P. Fairfax

**Author notes:** Equal contributions (Robert A. Watson, Chelsea A. Taylor).

## Abstract

Immune checkpoint blockade (ICB) has markedly improved outcomes across a range of tumours, including metastatic melanoma (MM). However, peripheral biomarkers of response remain lacking and underlying mechanisms of action incompletely described. A number of studies have demonstrated the value of T cell receptor (TCR) repertoire analysis in determining associations with response, however identifying key groups of T cells based on their TCR usage has remained elusive. Here we performed deep sequencing of the TCR of CD8^+^ T cells isolated from peripheral blood of patients receiving ICB for MM (n=91) at multiple time points, along with healthy control samples (n=42) and resected tumour specimens (from n=7 patients). Using the GLIPH2 algorithm to cluster TCR based on putative shared antigen specificity, we describe groups of TCR which expand post-ICB in responding patients which we term ‘Emergent Responder’ (ER) clones. We find that these ER clones are typically large and of a memory phenotype, with increased expression of genes encoding cytotoxic proteins. Analysis of tumours resected in advance of ICB demonstrates ER clones are enriched and expanded within the tumour compared to the periphery at pre-treatment. Significantly, we note the proportion of the peripheral repertoire occupied by ER clones strongly correlates with long-term clinical response. Clinical outcome further associated with HLA type and, crucially, can be validated across replication and independent datasets. This work provides the first-in-kind description of TCR-defined CD8^+^ T cells that mediate the response to ICB in MM, demonstrating the prognostic utility of the peripheral immune repertoire with potential widespread therapeutic and prognostic applications.

## INTRODUCTION

ICB works to re-invigorate the immune response to cancer and is sufficient to enable durable disease control in a significant subset of patients. This is most notable in MM, where combined blockade of CTLA-4 and PD-1 (cICB) is associated with >50% of patients surviving beyond five years (Larkin et al. 2019), the majority of whom were treatment free by this time-point (Regan et al. 2021). This disease control is primarily T cell mediated, with CD8^+^ T cells forming the primary effectors of ICB induced anti-tumour responses (A.C. Huang et al. 2017; Waldman, Fritz, and Lenardo 2020). It has long been recognised that features of the tumour-infiltrating (TI) T cell repertoire are associated with clinical responses to ICB, with increased clonality of this compartment predicting both overall survival (OS) and response to ICB across a range of cancer types (Zhang et al. 2020; Valpione, Mundra, et al. 2021; Yusko et al. 2019). Similarly, there is robust data to demonstrate TI T cells can also be observed in the peripheral blood (A.C. Huang et al. 2017) where they show comparatively reduced expression of exhaustion markers (Lucca et al. 2021; Wu et al. 2020). Treatment with ICB leads to mobilisation of peripheral clones (Yost et al. 2019), enhancing trafficking into the tumour and altering the relationship between these two pools. As such, peripheral blood can act as a ‘window’ into the TI compartment, providing on-treatment clinically-relevant insights (Kato et al. 2021; Zhang et al. 2020; Henick 2020). In keeping with this, peripheral CD8^+^ T cell repertoire features have correspondingly been correlated with response to ICB in MM and other tumours (Naidus et al. 2021; Dong et al. 2021; Kato et al. 2021; Cha et al. 2014; Valpione, Galvani, et al. 2020; Watson et al. 2021; Fairfax et al. 2020). However, whilst the association between broad repertoire features and a response to ICB is now established, a description of key CD8^+^ T cell clones based on their TCR sequence is currently lacking. Thus far there has been limited identification of neo-antigen specific TCR and, given most are likely patient-specific, the utility of such TCR to general therapeutics and prognostication is questionable. Conversely, despite melanomas having a high burden of private mutations, they demonstrate conserved patterns of gene expression. Within patients, the occurrence of specific common side-effects, notably vitiligo which associates with a favourable prognosis (Lommerts, Bekkenk, and Luiten 2021), is testament to likely shared T cell anti-tumoural mechanisms and receptor patterns.

We reasoned that by integrating a large dataset of CD8^+^ T cell TCR with long-term clinical outcome data we could identify shared features of TCR associated with durable clinical response to ICB in MM which may have both therapeutic and prognostic value. To do this we have used deep T Cell Receptor sequencing (TCRseq) of sorted peripheral CD8^+^ T cells from both healthy donors and a large cohort of patients undergoing ICB treatment for MM. By using the GLIPH2 algorithm (H. Huang et al. 2020) to cluster TCR based on putative shared antigen specificity, we describe groups of TCR which expand post-ICB in responding patients and hence term these ‘Emergent Responder’ (ER) clones. We find that these ER clones are typically large and of a memory phenotype, with increased expression of genes encoding cytotoxic proteins. Analysis of tumours resected in advance of ICB demonstrates ER clones are enriched and expanded within the tumour compared to the periphery at pre-treatment. Significantly, we note the proportion of the peripheral repertoire occupied by ER clones strongly correlates with long-term clinical outcome. We also find clinical outcomes associate with HLA-type and crucially, can be validated across replication and independent datasets, providing the first-in-kind description of TCR-defined CD8^+^ T cells that mediate the response to ICB in MM.

## RESULTS

### Identification of key response groups of CD8^+^ T cells

We performed deep sequencing of the TCR of sorted peripheral CD8^+^ T cells from 91 patients receiving ICB for MM, at multiple time-points before and after treatment, as well as from 42 healthy donors (Table S1). Using this dataset we replicated our previous observation that a greater number of large CD8^+^ T cells at 14-28 days post-treatment, defined as clones representing >0.5% of the total repertoire, associated with ongoing clinical responses six months post-treatment initiation ((Fairfax et al. 2020), Figure S1a).

Whilst we have previously demonstrated the presence of key clones in the response to MM in the peripheral blood prior to ICB (Watson et al. 2021), we were intrigued by the finding that the number of large clones at 14-28 days, but not at baseline, associated with outcome. We therefore hypothesised that we might be able to identify key clones which ‘emerge’ or expand following ICB. To investigate this further we used the GLIPH2 algorithm (H. Huang et al. 2020) to arrange TCRs into ‘specificity groups’ of putative shared antigen recognition. We filtered these using similar thresholds as previously described (see methods; (Chiou et al. 2021)), with a resultant 13,450 highly-public, expanded, specificity groups identified termed as ‘public specificity groups’ (Figure 1a).

**Figure 1.**
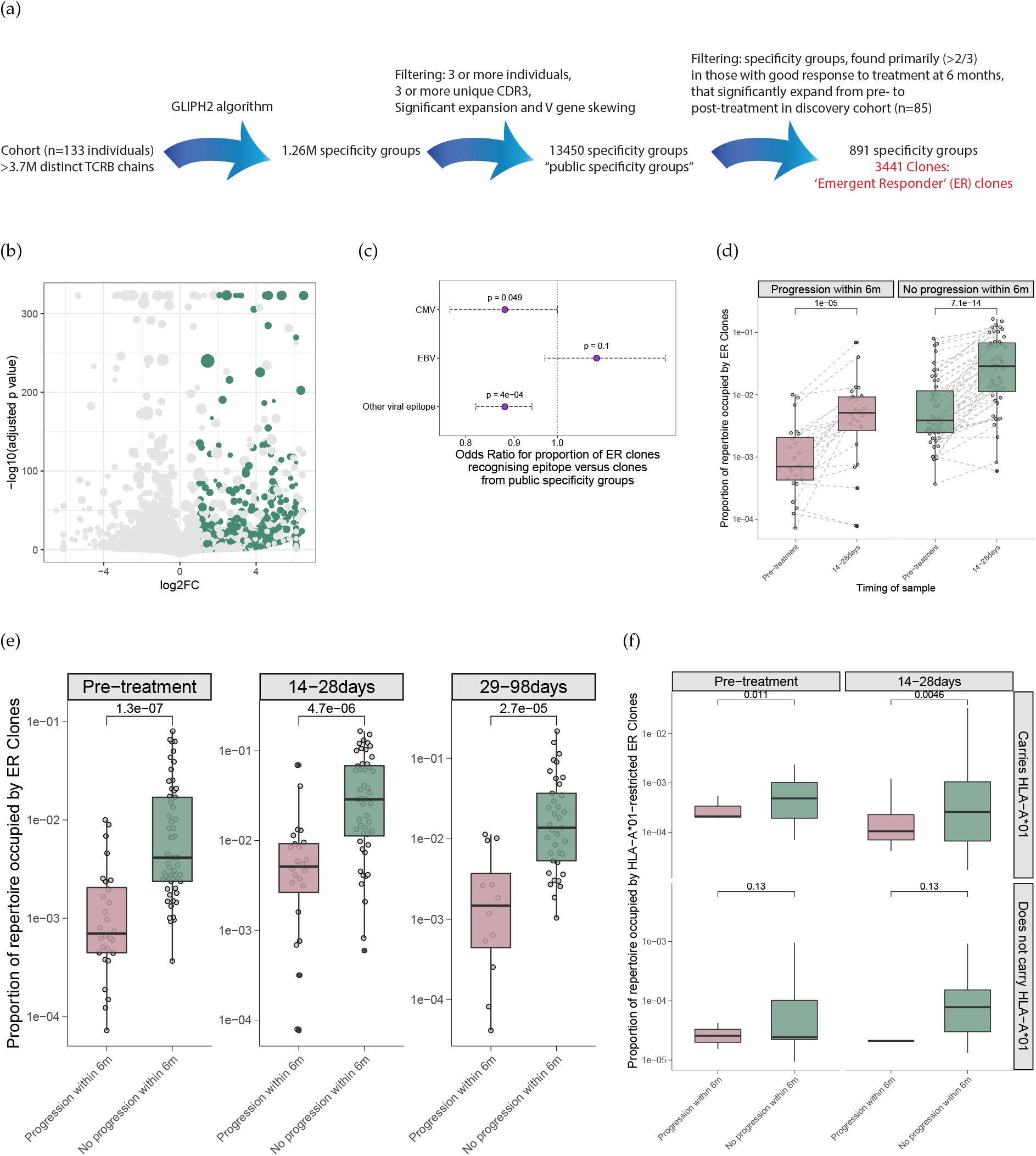
(a) Flow diagram depicted data filtering and post-processing of GLIPH2-clustered data. (b) Volcano plot showing change in size of filtered specificity groups pre- and post-treatment with ICB. ER specificity groups are coloured in green. (c) Odds ratio for putative antigen recognition of ER clones compared to recognition of the same antigen by clones from public specificity groups. (d) Proportion of repertoire occupied by ER clones coloured by outcome (disease progression at 6 months) and faceted by outcome. (e) As per (d) but faceted by time of sampling. (f) Proportion of repertoire occupied by HLA-A*01-restricted ER clones faceted by whether the individual carriers the HLA-A*01 allele. Statistics with Fisher’s exact test (c), Wilcoxon signed rank test (d) two-sided Wilcoxon rank-sum test (e) or one-sided Wilcoxon rank-sum test (f).

Within our cohort, 85 individuals had paired pre- and post-treatment blood samples (pretreatment samples taken immediately prior to the first dose of ICB, post-treatment samples taken a median of 21 days (IQR 7 days) following the first dose of ICB). These 170 samples were used as a “discovery cohort” to identify specificity groups, found primarily in those with a good response to treatment at six months, that significantly increased in size following ICB (see methods, Figure 1a). We found 891 specificity groups, comprising 3441 different clones, that minimally doubled in size and were primarily found in individuals with a positive response to ICB (Figure 1a and 1b). We term the clones within these specificity groups ‘<u>E</u>mergent <u>R</u>esponder’ (ER) clones i.e. those clones which are in emerging/enlarging groups post-treatment in individuals with a clinical response to ICB.

We found that there was no relationship between the proportion of the repertoire occupied by ER clones with either age or sex (Figures S1b, S1c). However, we did note a relationship with Cytomegalovirus (CMV) serostatus, with CMV positive patients having a proportionally lower repertoire occupancy by ER clones at all time-points (Figure S1d). Consistent with this, ER clones were not enriched for CMV or Epstein-Barr virus (EBV) clones as called by The Immune Epitope Database (IEDB) when compared to clones found in public specificity groups. Indeed, the point estimate suggests a trend towards depletion of clones recognising CMV with ER clones being significantly depleted for all viral epitopes (Figure 1c). In keeping with this, we found no association between CMV serostatus and clinical outcome (Figure S1e).

### ER clone repertoire occupancy corresponds with clinical response and is HLA specific

Having identified groups of TCR which significantly expand following ICB in those with a positive response to treatment (i.e. ER clones) we wished to verify that the presence of these key clones corresponded with clinical outcome. We found that the proportion of the repertoire occupied by ER clones significantly expanded with treatment in both ICB responders and non-responders (Figure 1d), although when we dichotomised by treatment type this expansion in non-responders was limited to recipients of cICB (Figure S2). However, surprisingly this was also observed across both pre-, as well as post-treatment time-points - individuals who responded to treatment demonstrated a significantly higher proportion of ER-clone repertoire occupancy (Figure 1e). This is in keeping with the baseline clonal repertoire being key to mediating ICB responses, consistent with previous findings (Watson et al. 2021).

Given specificity groups may demonstrate HLA type restriction (H. Huang et al. 2020), we explored the relationship between MM specificity groups and HLA class I type. We identified specificity groups associated with HLA allele carriage (Figure S3a) and thus were able to find corresponding HLA-specific ER clones. Notably, for several HLA types we were powered to observe outcome associations when ER clones were restricted to those that are members of specificity groups that significantly associate with an HLA allele. In HLA-A*01 carrying patients, this reduced the 3441 clones to just 68. Remarkably, the proportion of the repertoire occupied by these HLA-A*01-restricted ER clones associated with clinical outcome in both pre- and post-treatment samples, but restricted to individuals carrying an HLA-A*01 allele (Figure 1f). Likewise, we were able to repeat this observation for HLA-B*07 (71 clones) and HLA-C*05 (108 clones) restricted ER clones, two of the most common HLA types (Figures S3b and S3c).

### ER clones are large, persistent and of a memory phenotype

We have previously observed that the number of ‘large’ peripheral blood CD8^+^ T cell clones (defined as >0.5% of the repertoire) correlates with clinical response (Figure S1a, Fairfax et al. 2020), finding large clones show increased sensitivity to ICB (Watson et al. 2021). We therefore performed comparative analysis of ER clone size, finding that ER clones were more likely to be large clones (Figure 2a) with 2.8% of ER clones being large, compared to 2.2% of clones that are found within a public specificity group and 0.2% of all clones (OR 1.27, p=0.004 compared to clones in public specificity groups; OR 14.9, p<2.2×10^-16^ compared to all clones; Fisher’s exact test). Notably, there was a relationship between ER clone size and ICB response, with significantly more large clones also being ER clones in those who responded to ICB compared to those who progressed within six months (Figure 2b).

**Figure 2.**
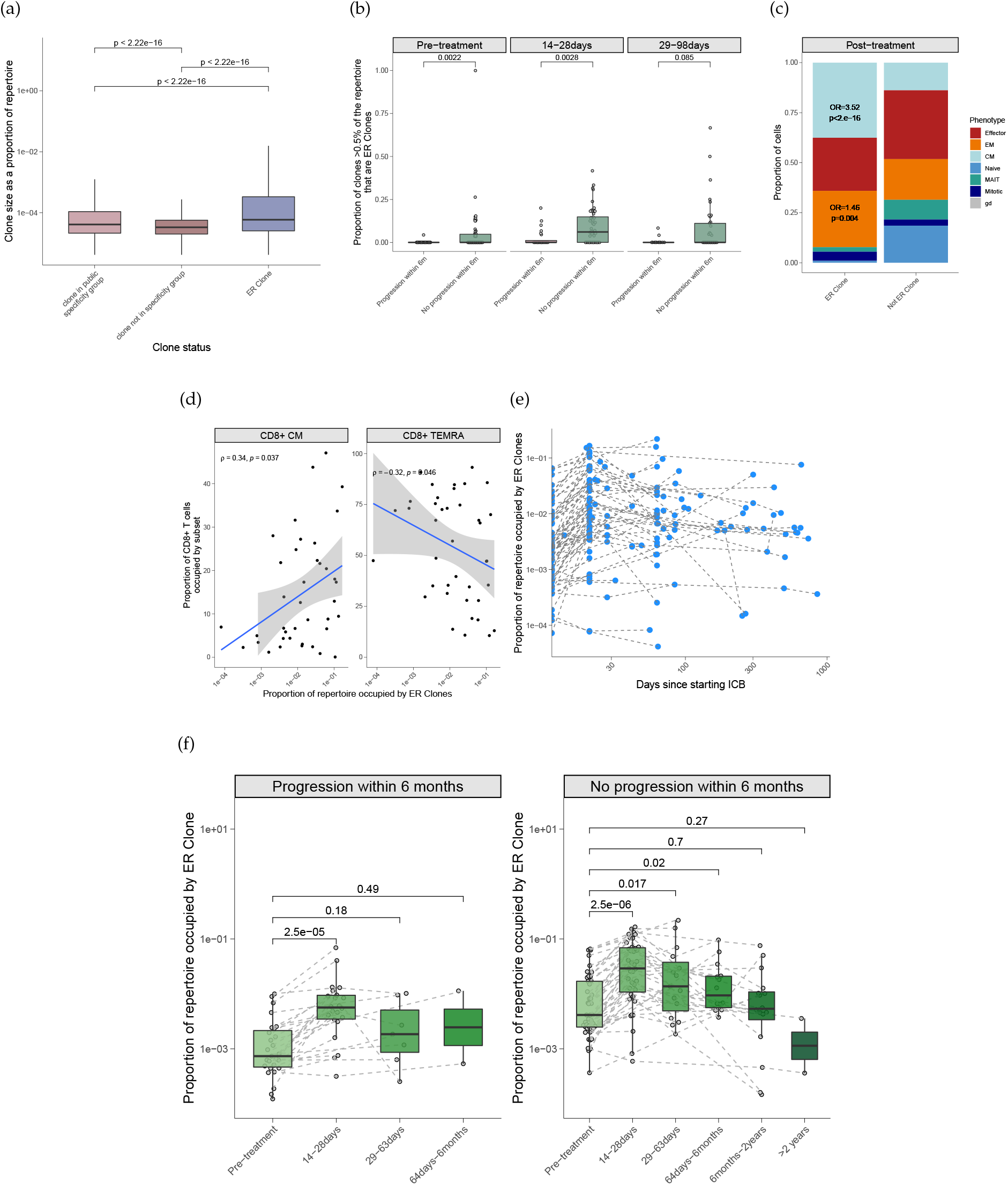
(a) Median size of ER clones as compared to clones from public specificity groups and all other clones. (b) Proportion of large clones (those occupying >0.5% of the repertoire) that are ER clones, split by clinical response and faceted by time point. (c) Proportion of cells by phenotype as defined by scRNAseq from (Watson et al. 2021), split by whether or not the cell is an ER clone - post-treatment time point. (d) Correlation between proportion of repertoire occupied by ER clones and central memory (left panel) or TEMRA (right panel) subsets as determined by flow cytometry at 14-28days post-treatment. (e) Proportion of the repertoire occupied by ER clones over time - each dot is an individual. (f) Same data as per (e) but samples grouped into time bins rather than continuous variable, and faceted by clinical response. Statistics are via two-sided Wilcoxon rank sum test (a, b and g), a Fisher’s exact test (c), Spearman’s rank correlation (d).

In order to examine the phenotypic subset of ER clones, we looked at single-cell RNA sequencing (scRNAseq) data of peripheral CD8^+^ T cells taken pre-treatment and 21 days post-treatment in patients receiving ICB for MM (n=4 patients receiving single-agent Immune Checkpoint Blockade; either Nivolumab or Pembrolizumab (sICB) and n=4 receiving combination Immune Checkpoint Blockade; Ipilimumab plus Nivolumab (cICB)) - as previously reported (Watson et al. 2021). We were able to identify 444 cells from 63 clones within this dataset that carry an ER clone ID (identical TRBV gene, TRBJ gene and complementary determining region 3 (CDR3) amino acid sequence). These cells were significantly enriched for both Effector Memory (EM) and Central Memory (CM) subsets both pre- and post-ICB (post-treatment OR 1.46, 95% CI 1.12-1.88, p=0.004 and OR 3.52, 95% CI 2.76-4.47, p<2.2×10^-16^, respectively, Fisher’s exact test) (Figures 2c). Notably, effector T cell (T_eff_) cells were under-represented in ER clones (OR 0.65, 95% CI 0.50-0.85, p=0.001).

To corroborate this, we performed flow cytometry data on PBMCs from a subset of samples (n=39), noting a positive correlation between the proportion of CD8^+^ T cells 14-28 days posttreatment that were CM on flow cytometry and the proportion of the repertoire occupied by ER clones (Figure 2d). There was an anti-correlation for CD8^+^ TEMRA cells, a surrogate for T_eff_ cells.

Based on the memory phenotype of ER clones, we then explored the long-term persistence of these clones in the peripheral blood. We looked at samples taken over two years following commencement of ICB and noted that the proportion of the repertoire occupied by ER clones was relatively stable over time (Figure 2e). In order to equate long-term changes with outcome, we grouped time-points into categories and faceted individuals by progression status at six months. Notably, whilst across both groups there is an initial increase in the proportion of the repertoire occupied by ER clones (as described above), this is short-lived for those with progression with ER clone repertoire occupancy returning to pre-treatment levels within 63 days. However, in patients without progression in the first six months of treatment, early expansion of ER clones is more marked with repertoire occupancy staying above pre-treatment levels for up to six months (Figure 2f).

### ER clones are enriched and expanded within tumours and are cytotoxic in the peripheral blood

In order to further investigate whether ER clones are tumour associated, we extracted RNA from ten tumour blocks (n=7 patients) from whom we had paired pre- and post-ICB peripheral blood data. Tumour specimens were resected a median of 6.4 months prior to the commencement of immunotherapy (range 3.5 weeks to 16.4 months). Remarkably, we found multiple ER clones within the tumours prior to commencement of ICB, in excess of that expected by chance based on the numbers present in peripheral blood (Figure 3a). Further, when restricting only to clones found in both the tumour and peripheral blood at baseline in the same patient, ER clones were significantly expanded in the tumour versus the peripheral blood when compared to clones of a similar size from public specificity groups (Figure 3b). Given the tumour blocks will have contained both CD4^+^ T cells and CD8^+^ T cells, the actual tumour-enrichment and expansion will be greater than inferred given the blood comparison is with the proportion of CD8^+^ T cells only.

**Figure 3.**
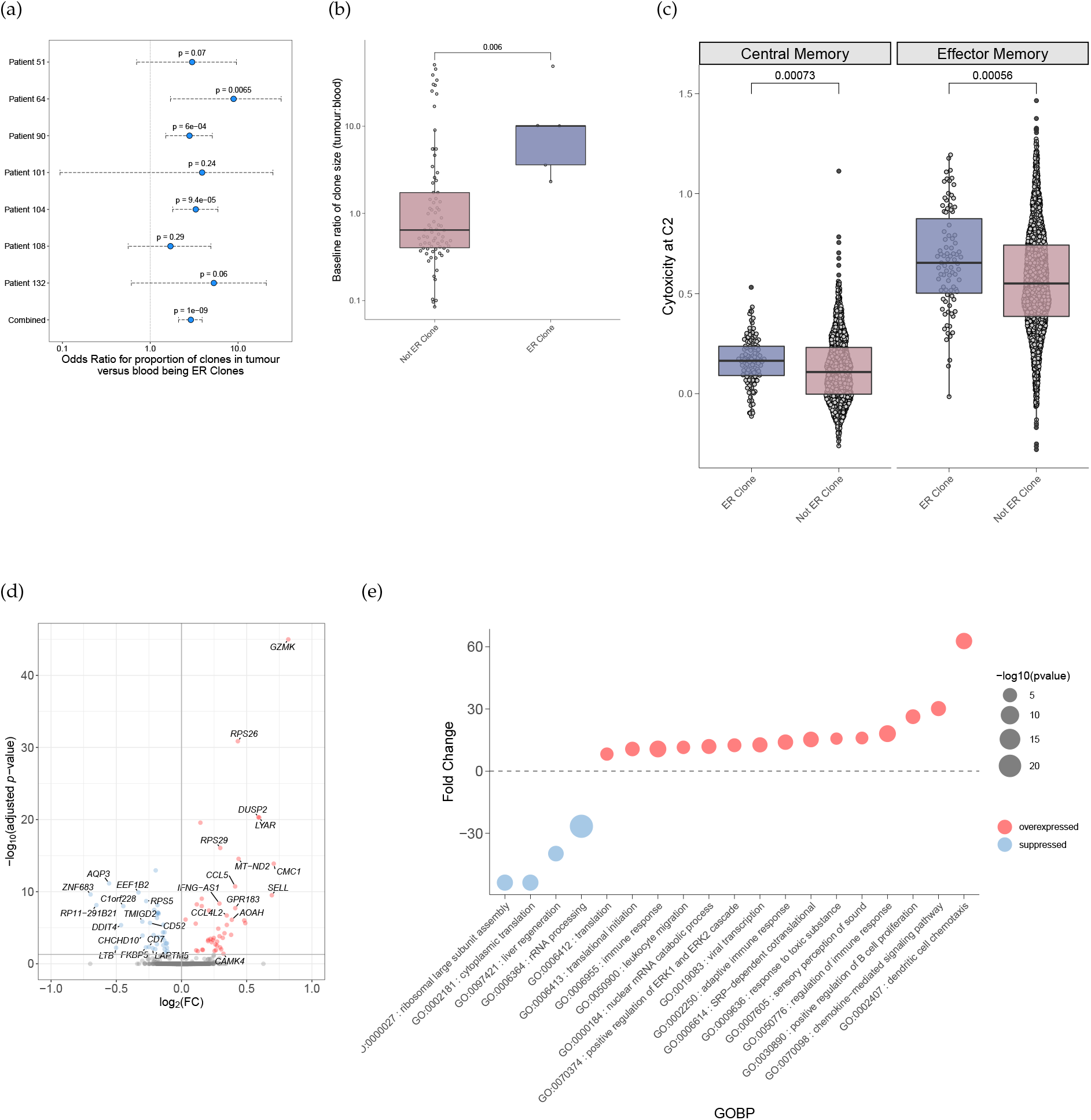
(a) Odds ratios for the proportion of clones within FFPE tumour specimens that are ER clones compared to the proportion of clones in the peripheral blood from the same patient. Whiskers show 95% CIs. Statistics with Fisher’s exact test. (b) Ratio of size of clone within tumour:size of clone within blood for clones that are found in both the peripheral blood and in FFPE tumour biopsies from the same patient, coloured by whether or not the clone is and ER clone. Statistics with one-sided Wilcoxon rank sum test. (c) Cytotoxicity score (based on previously described gene expression profile (Watson et al. 2021)) of cells coloured by ER clone, faceted by subset. (d) Volcano plot of genes upregulated (red) or downregulated (blue) in cells that are in ER clones versus those that are not. (e) Gene ontology biological process (GOBP) analysis for pathways which are overexpressed (red) or under-expressed (blue) in ER clones.

We then looked at the gene expression of those CD8^+^ T cells which are ER clones in a published scRNAseq dataset (Watson et al. 2021). Applying a previously-defined cytotoxicity score reveals that ER clones are more cytotoxic than non-ER clones within both central and effector memory T-cell clusters (Figure 3c). Specific genes differentially upregulated within ER clones include the central memory marker *SELL*, encoding CD62L, and *CCL5* encoding the cytokine of the same name, as well as cytotoxic molecules such as *GZMK* (Figure 3d, supplementary table S2). Gene ontology biological process (GOBP) pathway analysis reveals upregulation of adaptive immune pathways and translation, with downregulation of innate responses (Figure 3e).

Taken together, these findings suggest increased tumour-reactivity of ER clones which, in the peripheral blood, are more cytotoxic with upregulation of key immune pathways, including those involved in chemotaxis and tissue trafficking.

### ER clones are found within the repertoires of healthy donors

ER clones are defined on the basis of being highly-public i.e. found in at least three different individuals. We therefore wished to understand their occurrence within the T cell repertoires of healthy donors. Surprisingly, ER clones are found within healthy donor repertoires, although they occupy a smaller proportion of the repertoire to patients with MM pre-treatment (median proportion in patients pre-treatment 0.26%, median proportion in healthy donors 0.057%, p<0.0001, Figure 4a). This is due to the fact that patients with MM have both more and larger ER clones (Figures S4b and S4a). However, when patients were split by whether they respond to ICB within the first six months of treatment, we noted that only patients with disease control for over six months had a greater proportion of their repertoire occupied by ER clones than healthy donors (HDs) (Figure 4b). Those individuals with progressive disease had a similar proportion of their repertoire occupied by ER clones compared to HDs. These results suggest that failure to respond to ICB could be a result of a lack of expansion of key CD8^+^ T cell clones at baseline, indicating the proportion of repertoire occupied by ER clone at baseline may have prognostic utility.

**Figure 4.**
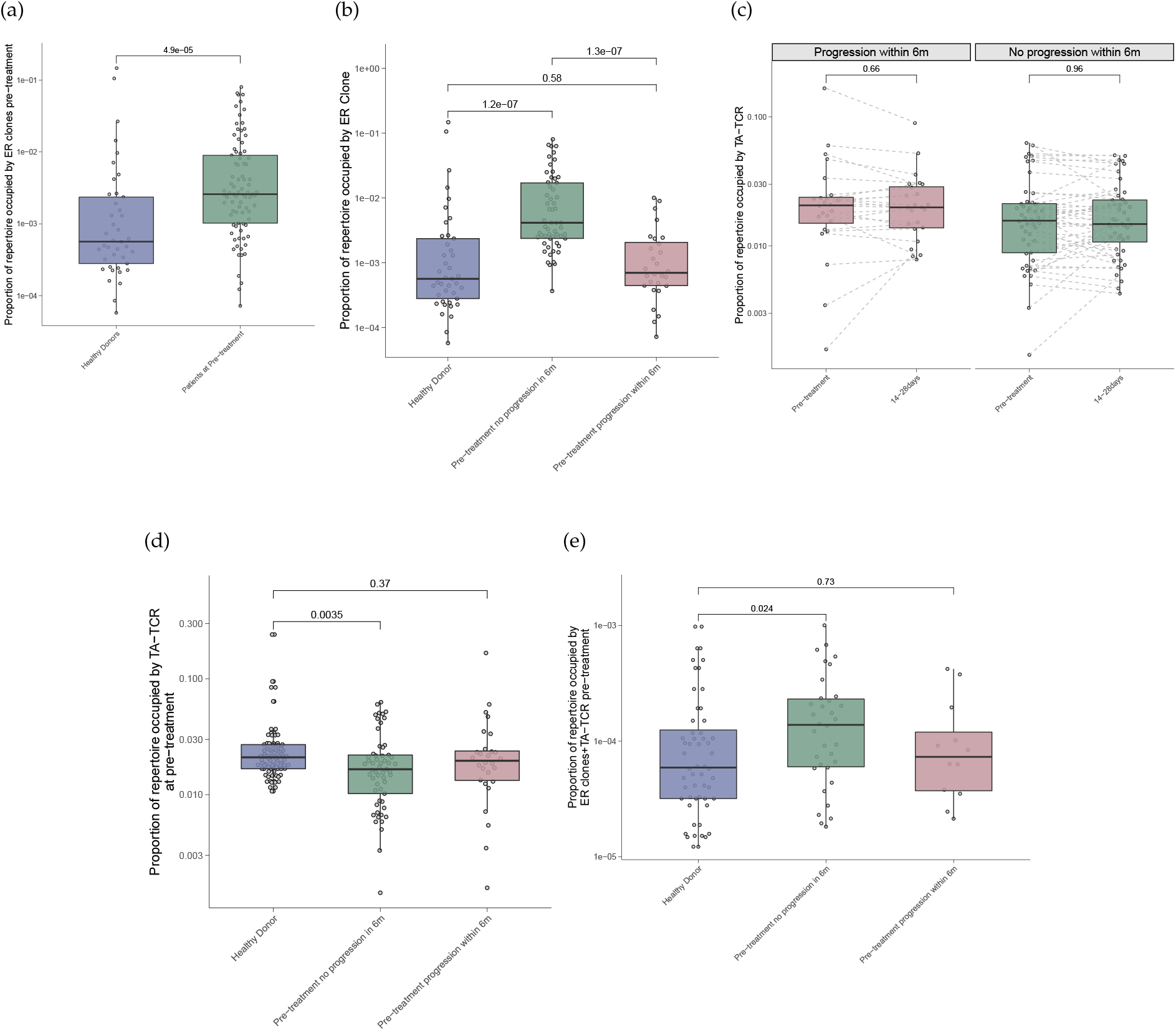
(a) Proportion of the repertoire occupied by ER clones in healthy donors and patients pre-treatment. (b) Repertoire occupancy by ER clones comparing healthy donors to patients with no progression of disease in 6 months (green) and progression of disease within 6 months (red). (c) Proportion of the repertoire occupied by TA-TCR by clinical response over time. (b) Repertoire occupancy by TA-TCR comparing healthy donors to patients with no progression of disease in 6 months (green) and progression of disease within 6 months (red). (c) as per (b) but for TA-TCR that are also ER clones (n=91). Statistics with two sided Wilcoxon rank sum test (a, b, d and e) or Wilcoxon signed rank test (c).

For comparison, we took known tumour-associated TCR (TA-TCR) which had been derived from sequencing of TCRβ chains extracted from resected melanomas, within published literature (Riaz et al. 2017; Yusko et al. 2019; Tumeh et al. 2014; Pruessmann et al. 2020). Within our dataset, 20,553 unique TA-TCR were present. However, unlike ER clones, there is no effect of treatment on TA-TCR clones, with no evidence of expansion post-ICB in either those individuals who respond or progress (Figure 4c). Further, we noted that the repertoire occupancy of TA-TCR is similar between patients with MM pre-treatment and healthy donors with those who have a good response actually having less of their repertoire occupied by TA-TCR than HDs (Figure 4d). Finally, we looked within the TA-TCR found in our dataset and identified that 91 are also ER clones. These clones, which are both TA-TCR and ER clones, are expanded at baseline in individuals with a response to treatment akin to all ER clones but in contrast to the rest of the TA-TCRs (Figure 4e).

Taken together, these findings suggest that most TA-TCR are not expanding with ICB and therefore may not be being ligated by antigen. Indeed, it is plausible and in keeping with other observations that most TA-TCR are so-called “bystander clones” whereas, in contrast, those identified as being ER clones are, in fact, mediating an anti-tumour response (Scheper et al. 2019).

### Importance of ER clones replicates in validation datasets

Finally, we sought to validate the relevance and importance of ER clones in external datasets. In order to replicate the observation that the proportion of the peripheral blood CD8^+^ T cell repertoire occupied by ER clones correlated with outcome, we performed bulk RNA sequencing on sorted peripheral CD8^+^ T cells from a further 95 individuals (i.e. not included in the discovery cohort) with MM (peripheral blood validation cohort, n=53 individuals paired pre- and post-ICB, Table S3). V(D)J data was mapped from bulk RNA sequencing (bulk RNAseq) fastq files using MiXCR (Bolotin et al. 2015) hence this dataset of separate individuals comprised TCR repertoire data generated using an alternative method.

Within this dataset, we identified 139 unique ER clones, with at least one clone being found in 73/95 individuals. We were able to replicate the finding that the proportion of repertoire occupied by ER clones associated with outcome both at pre-treatment and post-treatment time points (Figure 5a). Furthermore, the proportion of the repertoire occupied by ER clones in the peripheral blood at both the pre-treatment and 14-28 day time-point resulted in both a progression-free and overall survival benefit (Figure S5). After confining analysis to the validation cohort, individuals with greater than the median proportion of their repertoire occupied by ER clones at 14-28 days were 63% less likely to progress within six months and 56% less likely to die than those with less than the median of their repertoire occupied by ER clones after controlling for age (HR for progression 0.37, 95% CI 0.16-0.83, p=0.016; HR for death 0.44, 95% CI 0.20-0.98, p=0.045; Cox proportional hazards). Using the bulk RNAseq data and including all individuals (including those from the discovery cohort (n=178 total, n=170 pre-treatment and n=131 14-28 days post-treatment)), there was a highly significant association between proportion of the repertoire occupied by ER clones, both pre- and post-treatment, and clinical outcome (Figures 5b, 5c, and S5).

**Figure 5.**
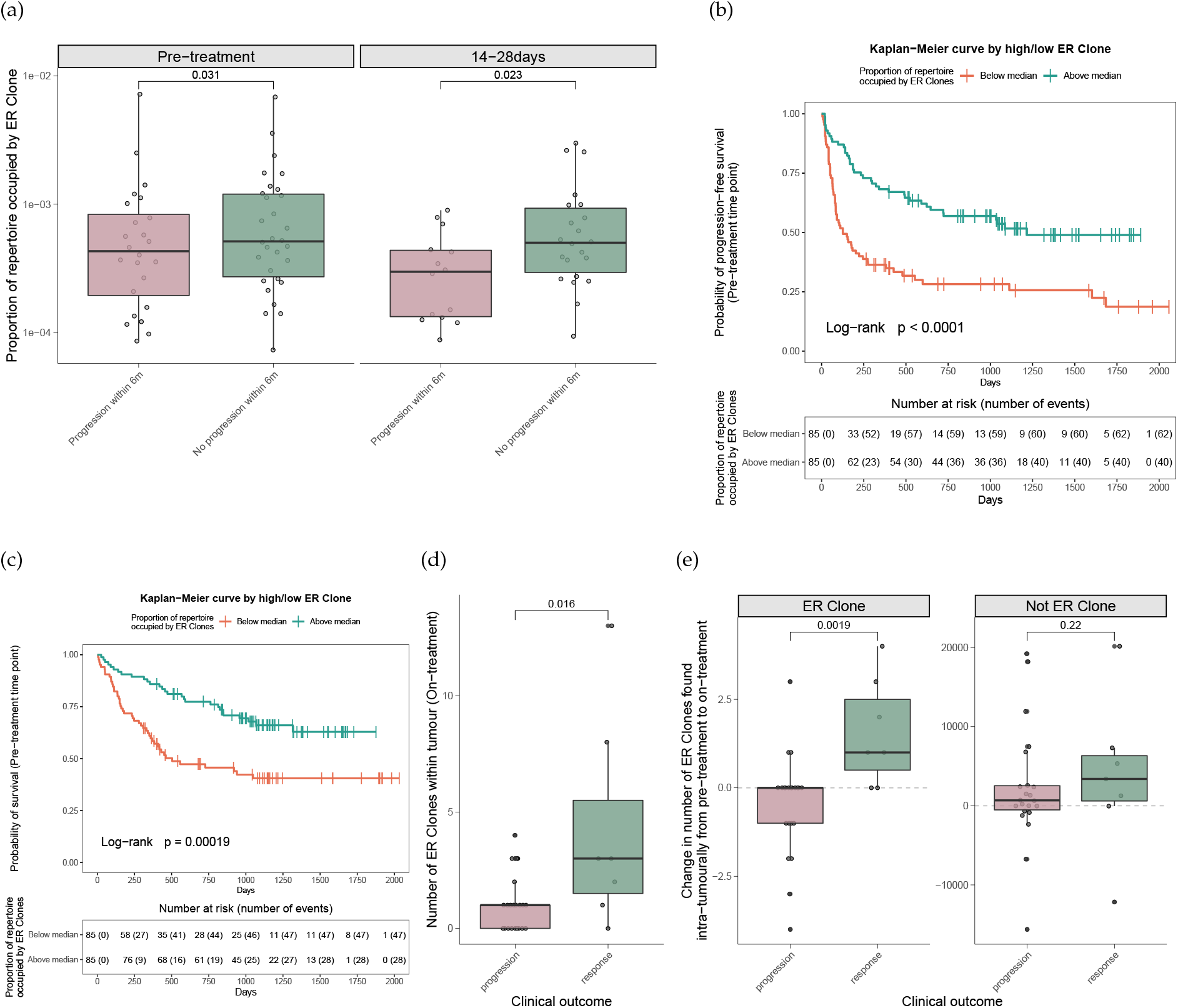
(a) Proportion of repertoire occupied by ER clones in validation cohort (Fairfax MiXCR) coloured by whether there was progression of disease within the first six months of treatment, faceted by timing of sample. (b) Kaplan-Meier curve for progression-free survival based on above/below median proportion of repertoire occupied by ER clones pre-treatment in pooled bulk RNAseq cohort (patients from validation plus discovery cohort). (c) as per (b) but for overall survival. (d) Number of ER clones found within tumours at the on-treatment time point from Riaz et al (Riaz et al. 2017) on-treatment, coloured by clinical response. (e) Change in the number of clones found within tumours, coloured by clinical outcome and faceted by whether or not clones are ER. Data taken from same dataset as (d). Statistics with one-sided Wilcoxon rank sum (a, d and e) and Log rank test b and c.

In order to corroborate our findings of the intra-tumoural relevance of these clones, we explored the dataset from Riaz et al (Riaz et al. 2017) who sequenced TCR from resected melanomas pre- and post-sICB therapy. We find that the number of ER clones found within the tumours in post-treatment samples correlated with outcome in this study too (Figure 5d). Further, there appeared to be trafficking of ER clones into the tumours in patients that responded to PD-1 blockade, but not in those with progressive disease (Figure 5e). Such a difference was not observed in non-ER clones.

These results suggest that ER clones as defined in the discovery cohort are highly-public and of broad disease relevance across multiple datasets in MM.

## DISCUSSION

Here we use deep sequencing of the TCR of peripheral CD8^+^ T cells, coupled with unsupervised clustering using the GLIPH2 algorithm (H. Huang et al. 2020), to identify highly-public T cell clones which strongly associate with response to ICB treatment in metastatic melanoma. These clones, termed ‘Emergent Responder’ (ER) clones, are large, cytotoxic and of a memory phenotype. They show a tendency to upregulate tissue-trafficking chemokines and are found to be enriched and enlarged within tumours. Further, ER clones persist long-term in the peripheral blood, with durability of this expansion being associated with a sustained response.

This approach describes, for the first time, TCR from peripheral CD8^+^ T cells which strongly correlate with outcome in MM, building on previous work from ourselves and others which has described more broad peripheral repertoire features of response (Cha et al. 2014; Kato et al. 2021; Fairfax et al. 2020; Valpione, Galvani, et al. 2020; Watson et al. 2021) and as such provides avenues for further work and investigation.

Firstly, these TCR have clear potential as a peripheral host immune biomarker for response to ICB. The fact that their presence in the repertoire corresponds with clinical outcome in both the discovery and validation datasets at multiple time points suggests that they could be used following either pre- or on-treatment blood sampling to stratify patient response.

Secondly, this work provides further mechanistic insight into the importance of the *a priori* CD8^+^ T cell repertoire in the response to ICB in MM. The fact that low numbers of these key TCR pre-treatment associates with poor outcomes suggests that one mechanism of treatment failure is insufficient antigen recognition, consistent with previous findings (Cha et al. 2014; Watson et al. 2021). Further, our finding that many of these key TCR are found in healthy individuals suggests that key response T cells may be cross-reactive and not restricted to private neo-antigens, as has been suggested for non-small cell lung cancer (NSCLC) (Chiou et al. 2021). Given that those with disease progression had a lower proportion of their repertoire occupied by ER clones than healthy donors suggests a role for a failure of general immune surveillance in those with a poor response to treatment.

Thirdly, for both cancer vaccines and autologous cell therapies, the identification of public neoantigens - tumour targets relevant to multiple individuals - is of great importance for both logistical and clinical reasons (Pearlman et al. 2021). By identifying highly public groups of key response TCR in an unsupervised manner, there is great potential to subsequently advance this current work and identify cognate antigens with possible import in these therapeutic fields. Our work indicates that responsivity in terms of change in clonal size to ICB is vital in determining public clonotypes with prognostic and potentially therapeutic utility - the absence of response to ICB of clonotypes corresponding to public TA-TCR, as well as the lack of prognostic association of such clones, indicates that focusing on tumours in isolation at a single time point is likely to be of relatively low return. Moreover, our data are in keeping with the majority of clones within tumours forming bystanders - the relative depletion of such clones in the blood of responding individuals potentially indicating that these may be non-benign in effect.

A recent study (Sidhom et al. 2022) has also attempted to describe TCR motifs associated with a successful response to ICB. Using a deep learning approach, this work notably described motifs associated with viral antigens corresponding with a higher likelihood of response, whilst tumour-specific TCR corresponded with a higher likelihood of non-response. This finding is in contradiction to our description of ER clones which are depleted of viral-recognising TCR. However, Sidhom *et al*’s study is based on the intra-tumoural repertoire, not the peripheral blood, and they describe that the viral signature that associates with response may be relative to generalised background TCR. The approach described here - which agonistically identifies groups of TCR expanding post-treatment in people with a good response to treatment - suggests that, despite being highly public, key response TCR appear to be depleted for virally-reactive clones and instead show enrichment for recognition of certain tumour antigens.

We recognise the limitations of this work. Firstly, the study design primarily allows for associations to be described, rather than causality to be assigned. Secondly, as with most studies of T cell repertoires using bulk data, we are limited by our ability to pair TCR beta and alpha chains and therefore identify the entire TCR. Thirdly, as is standard practice for all RNA-based TCRseq, there is an underlying assumption that number of reads correlates closely with clonal size. However, given that we have used multiple modalities including bulk sequencing, single cell sequencing and flow cytometry and that our findings validate in multiple external datasets using differing samples and technologies, we are confident in the robustness, relevance and high applicability of our findings.

In conclusion, we can identify highly-public groups of key peripheral CD8^+^ TCR which closely correspond with clinical outcome following ICB treatment for MM across multiple datasets. These TCR are present in the baseline and clones carrying them are expanded, more cytotoxic, tumour-reactive and of a memory phenotype. This work is of broad relevance to biomarker development and targeted immunotherapeutic development.

## Acknowledgements

We are very grateful to all patients who contributed samples and participated in the study. We thank all the staff of the Day Treatment Unit, Oxford Cancer Centre, and The Brodey Centre at the Horton General Hospital. We are grateful to all the staff of the Oxford University Hospitals NHS Foundation Trust - particularly Miranda Payne, Nick Coupe and Rubeta Matin in the cancer centre who aid with patient recruitment, as well as the staff of the Oxford Radcliffe Biobank (ORB) and Churchill Hospital Sample Handling Lab.

## Author contributions

RAW, CAT, BPF conceived the study. RAW, WY, BPF and MM recruited patients and annotated clinical outcome data. RAW, CAT, OT, RC, AVA, PKS and FMS collected blood samples. RC and EI identified and selected FFPE tumour specimens, RC extracted RNA from FFPE tumour specimens. RAW and CAT performed the TCRseq experiments. RAW, CAT, IN performed preprocessing of TCRseq. CAT, OT, AVA performed bulk RNAseq; IN performed pre-processing. OT, WY, AVA performed genotyping and pre-processing; EJ and JJG performed HLA-typing of genotyped data. CAT performed flow cytometry. RAW and RC performed scRNAseq experiments with RAW and OT performing downstream analysis of this data. RAW, CAT, BS, BSu performed analysis of repertoire features. RAW performed GLIPH analysis. RAW produced all figures. RAW and BPF drafted the manuscript. BPF oversaw the project.

## Data availability

Raw sequencing data will be made publicly available via EGA.

## Methods

### Participants

Adult patients referred to receive ICB as standard of care therapy for the treatment of MM at Oxford University Hospitals NHS Foundation Trust (OUHFT) were prospectively and consecutively approached for recruitment. All patients provided written informed consent to donate samples to the Oxford Radcliffe Biobank (ORB) (Oxford Centre for Histopathology Research ethical approval reference 19/SC/0173, project nos. 16/A019, 18/A064, 19/A114) and grant access to clinical data. Patients received either cICB (ipilimumab 3 mg/kg plus nivolumab 1 mg/kg 3 weekly for ≤4 treatment cycles, followed by maintenance nivolumab) or sICB consisting of either nivolumab 480mg monthly, pembrolizumab 2 mg/kg three weekly or pembrolizumab 4mg/kg six weekly. Patient characteristics are given in Tables S1 and S3. Healthy donor participants were recruited via the Oxford Biobank (www.oxfordbiobank.org.uk; ethical approval reference 06/Q1605/55), with written informed consent. All donors were of European ancestry, 104 were female, 66 male, aged between 21 and 66 years (median, 46.5 years; IQR 18 years).

### Clinical outcome data

Patient demographic and clinical characteristics were collected from the electronic patient record (EPR). Sex was determined on the self-reported assignment extracted from the EPR. Response data was obtained from the EPR with progression being defined either clinically or using radiological assessment according to irRECIST.1.1., performed approximately 12 and 24 weeks post-initiation of treatment. All patients who had received a minimum of one cycle of ICB were incorporated into the analysis irrespective of clinical outcome.

### Sample collection

30-50 mL blood was collected into EDTA tubes (BD vacutainer system) immediately prior to administration of ICB. Peripheral blood mononuclear cells and plasma were immediately obtained from whole blood by density centrifugation (Ficoll Paque). All cell subset isolation for RNA sequencing was carried out by magnetic separation (Miltenyi) using CD8 positive selection for CD8 T cells, with all steps performed either at 4°C or on ice.

### RNA extraction

Post-selection cells were resuspended in 350 μL of RLTplus buffer supplemented with betamercaptoethanol (BME) or Dithiothreitol (DTT) (final concentration 143 mM BME/40 mM DTT). Samples were snap-frozen at 80°C for batched RNA extraction. Homogenisation of the sample was carried out using a QIAshredder (Qiagen), followed by RNA extraction using the AllPrep DNA/RNA/miRNA kit (Qiagen). RNA was eluted into 34 μL of RNase-free water with concentration quantified by Qubit. DNA was eluted into 54 μl elution buffer with concentration determined by NanoDrop. Both RNA and DNA samples were subsequently stored at −80°C until preparation for sequencing and genotyping.

For FFPE tumour samples, blocks were identified using clinical histopathological reports. Haematoxylin and Eosin (HE) slides for each block were microscopically inspected to identify areas of tumour deposit. Blocks were sectioned into 10 micron slices in a clean and RNase-free environment. Unstained sections were macro-dissected to obtain tumour regions. Five unstained sections were then combined and deparaffinised prior to RNA extraction using the AllPrep DNA/RNA FFPE extraction kit (Qiagen) according to manufacturer’s instructions. RNA was eluted in 14 or 20 μl RNAase-free water.

### HLA-typing

Genotyping was performed on the Illumina Global Screening Array 24 v3 (Illumina). The vcf file exclusively containing chromosome 6 was used as an input for the Michigan Imputation Server Genotype Imputation HLA (Minimac4) 1.5.8 pipeline against the Multi-ethnic HLA reference panel. Results were filtered by Class I and II genes.

### TCR sequencing

TCR sequencing was performed using the QIAseq Immune Repertoire RNA Library Kit (Qiagen) according to manufacturer’s instructions. In short, RNA (median input 80ng) was reverse transcribed to cDNA before being adapter-ligated and amplified. Sequencing was performed on an Illumina Novaseq (SP flow cell). Alignment and mapping of FASTQ files was performed using the CLC genomics workbench (Qiagen) for immune repertoires using default settings. Mapped sequences were filtered for productive beta chains. A TCR clone was defined throughout as a concatenation of the V gene, J gene and amino acid sequence of the CDR3 region.

### GLIPH2 analysis

GLIPH2 was run according to package instructions with default parameters; the reference files were as provided (http://50.255.35.37:8080/). Results were filtered as described elsewhere (Chiou et al. 2021) - specificity groups found in 3 or more individuals, containing 3 or more clones and with significant biases in Vb gene usage, and CDR3b clonal expansion in comparison with the reference dataset.

To identify ER clones, we compared abundance (as a normalized count) of all TCR within each specificity group pre- and post-ICB, generating a p value for the comparison using the poisson.test function in R (alternative = ‘two-sided’) - as previously described (Chiou et al. 2021). Specificity groups were called as being ‘responder’ if >66% of individuals in which they were found did not have progression of disease within 6 months of starting treatment. Responder specificity groups were called as ER clones if the Log2FC from pre-to post-ICB was >1 with an FDR-adjusted p value of < 0.05.

### Single cell RNA sequencing

Data analysed was that which has been previously published (Watson et al. 2021). All methods describing the preparation and sequencing of samples, as well as access to all data, is available via this reference (https://doi.org/10.1126/sciimmunol.abj8825).

### Flow cytometry

Patient PBMCs frozen in 90% FCS + 10% DMSO were thawed at 37°C and washed once in HBSS prior to staining of 1×10^6^ cells with LIVE/DEAD Fixable Near-IR (Invitrogen). Cells were washed and subsequently stained with antibodies against CD3, CD4, CD8, CD27 and CD45RA in 5% FBS + HBSS. After washing, cells were fixed for 10 minutes with 2% paraformaldehyde (Sigma), washed and resuspended in 5% FBS + HBSS prior to acquisition on a BD LSRII. Data was analysed in FlowJo (Treestar, v10.7.1) and R (v4.0.5). All staining steps were performed for 30 minutes and at 4°C unless otherwise stated.

### Antigen-specific TCR calling

CDR3β chains were matched to epitopes using IEDB TCRMatch tool (http://tools.iedb.org/tcrmatch/) with the highest scoring epitope match being assigned to each CDR3β chain with a minimum score of 0.90 (IEDB ‘medium’ confidence) used as a lower cut-off.

### Tumour-associated TCR (TA-TCR)

Sequences and sample metadata were downloaded from published datasets (Riaz et al. 2017; Yusko et al. 2019; Tumeh et al. 2014; Pruessmann et al. 2020). Chains were filtered for TRB with resolved information on V gene, J genes and CDR3 amino acid sequence. Clones were defined based on a concatenation of the V gene, J gene and CDR3 amino acid sequence.

### Bulk RNA sequencing

Bulk RNA-sequencing was performed on CD8^+^ T cell samples derived from patients receiving single or combination ICB, as detailed above. RNA was thawed on ice prior to mRNA isolation using NEBNext Poly(A) mRNA Magnetic Isolation Module kits. Up to 600ng of RNA was then used to generate dsDNA libraries using NEBNext Ultra II Directional RNA Library Prep Kits for Illumina (cohort sizes given in the text) as previously described (Fairfax et al. 2020). Samples were then sequenced on either an Illumina HiSeq4000 (75bp paired-end) or a NovaSeq6000 (150bp paired-end). TCR analysis was performed using the MiXCR package (Bolotin et al. 2015) with settings as previously described (Fairfax et al. 2020; Watson et al. 2021).

### Validation in external intratumoural dataset

Sequences and sample metadata were downloaded from (Riaz et al. 2017). Clones were defined based on a concatenation of TRB V gene, J gene and CDR3 amino acid sequence. The number of clones that we identified as ER clones found in each tumour sample were counted, with response defined as per the original publication metadata (response = CR/PR, progression = PD).

### General statistical analysis

Statistical comparisons were performed as indicated in each figure legend and conducted in R (v4.0.5).

## Supplemental information

### Supplemental Tables

**Table S1.** Cohort characteristics. Numbers given are number of samples

| Characteristic | N = 289<sup>†</sup> |
| --- | --- |
| Donor Type |  |
| Healthy volunteer (n=42 individuals) | 42 (15%) |
| Patient (n=91 individuals) | 247 (85%) |
| Age | 64 (15.3) |
| Sex is Female | 131 (45%) |
| Treatment |  |
| cICB | 112 (45%) |
| sICB | 135 (55%) |
| Timing of sample |  |
| Pre-treatment | 88 (36%) |
| 14-28days | 74 (30%) |
| 29-98days | 52 (21%) |
| 99-365days | 16 (6.5%) |
| >365days | 17 (6.9%) |
| <sup>†</sup> n (%); Mean (SD) |  |

Table S2. Significant genes (adjusted p value < 0.05 differentially regulated in ER clones. *Provided as csv file*.

**Table S3.** Characteristics of validation cohort. Numbers given are number of samples with 98 individuals in total including 51 paired pre- and post-ICB samples

| Characteristic | N = 148<sup>†</sup> |
| --- | --- |
| Age | 62 (14.6) |
| Sex |  |
| Female | 73 (49%) |
| Male | 75 (51%) |
| Treatment |  |
| cICB | 83 (56%) |
| sICB | 65 (44%) |
| Timing of sample |  |
| Pre-treatment | 88 (59%) |
| 14-28days | 60 (41%) |
| <sup>†</sup> Mean (SD); n (%) |  |

### Supplemental Figures

**Figure S1.**
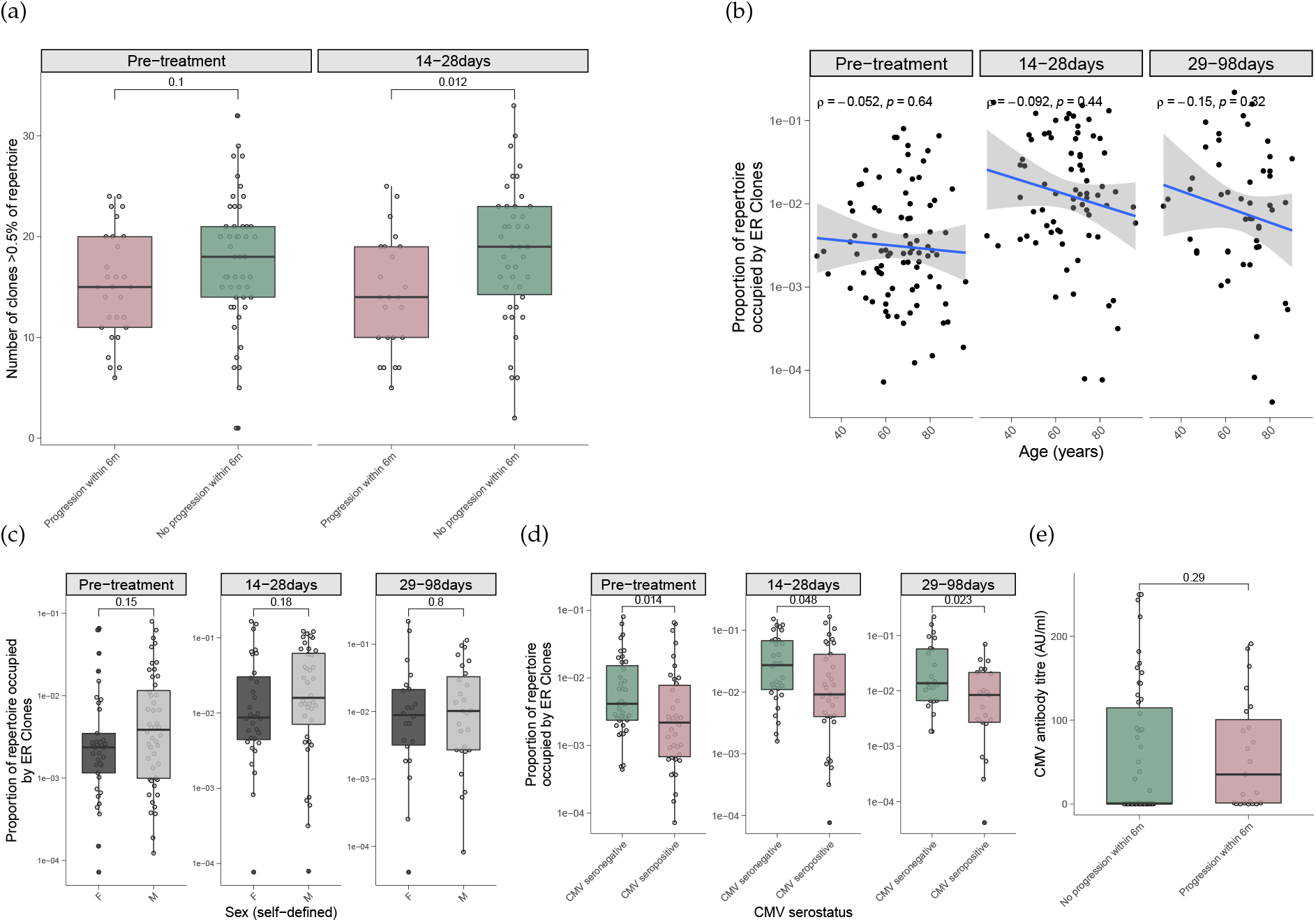
(a) The number of clones >0.5% of the repertoire, as defined by deep-sequencing, dichotomised by disease status six months post-treatment initiation. (b) Proportion of repertoire occupied by ER clones by age at pre- and post-treatment time-points. (c) Proportion of repertoire occupied by ER clones across time-points according to self-defined sex. (d) Proportion of repertoire occupied by ER clones across time-points according to CMV serostatus. (e) CMV titre coloured by clinical outcome. Statistics with Wilcoxon rank sum test (a, c-e) and Spearman’s rank correlation (b).

**Figure S2.**
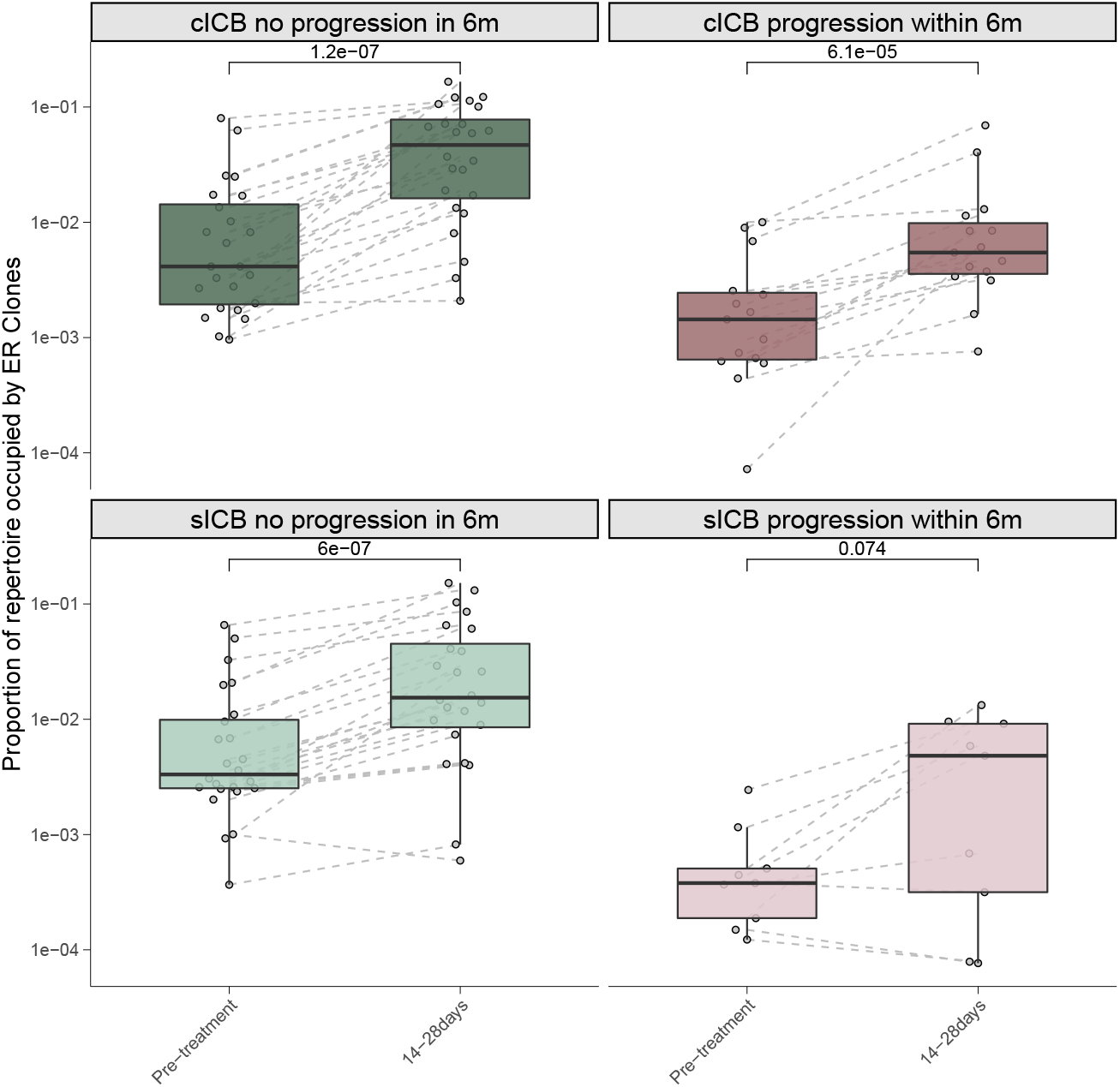
Proportion of the repertoire occupied by ER clones pre-treatment and at 14-28days faceted by treatment and clinical outcome. Statistics with Wilcoxon signed-rank test.

**Figure S3.**
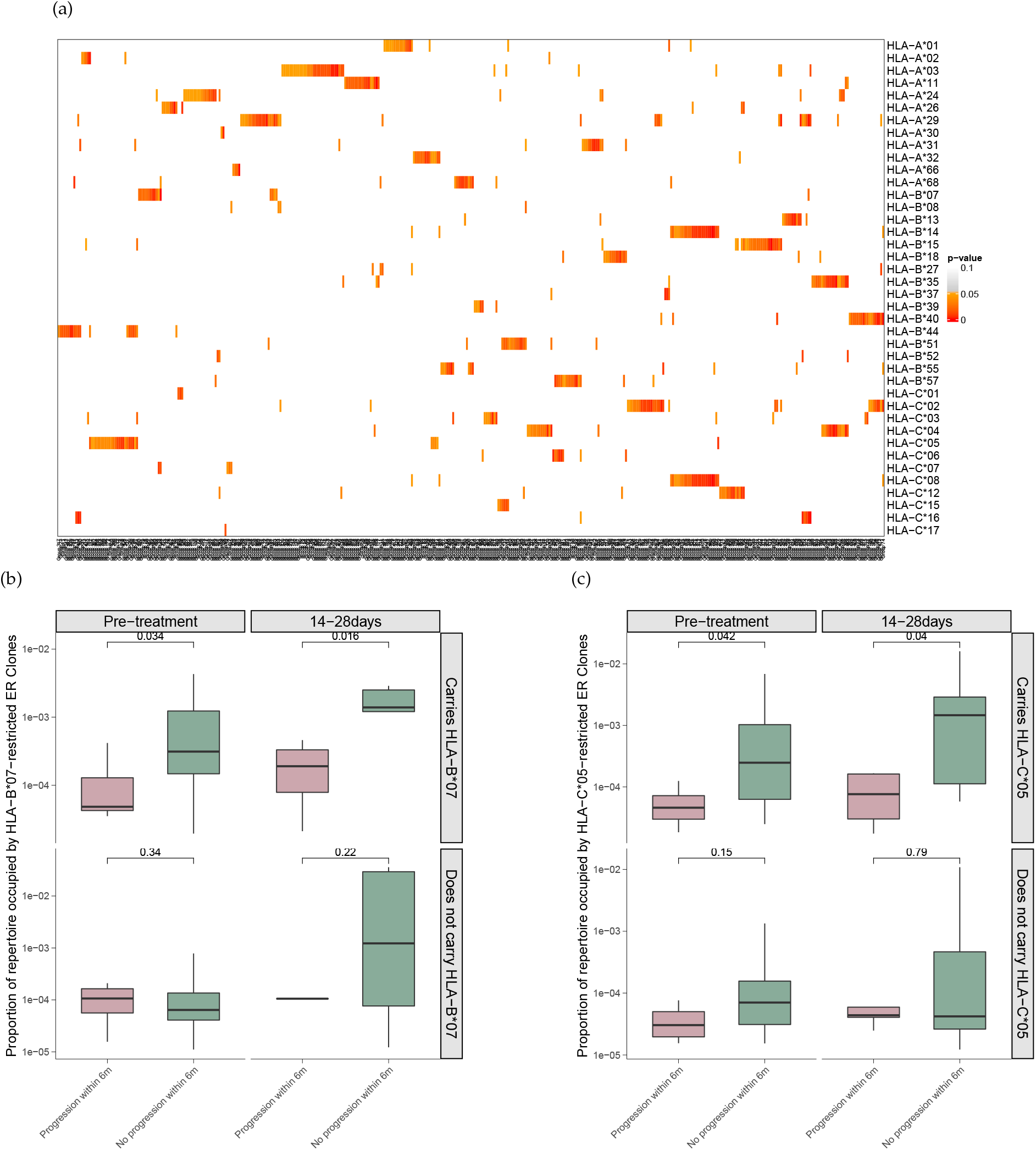
(a) Heatmap showing associations between ER clones (columns) and HLA class I type. (b) Proportion of repertoire occupied by HLA-B*07-restricted ER clones faceted by whether the individual carriers the HLA-A*01 allele. (c) As per (b) but for HLA-C*05. Statistics with Fisher’s exact test (a) or one-sided Wilcoxon signed rank test (b-c).

**Figure S4.**
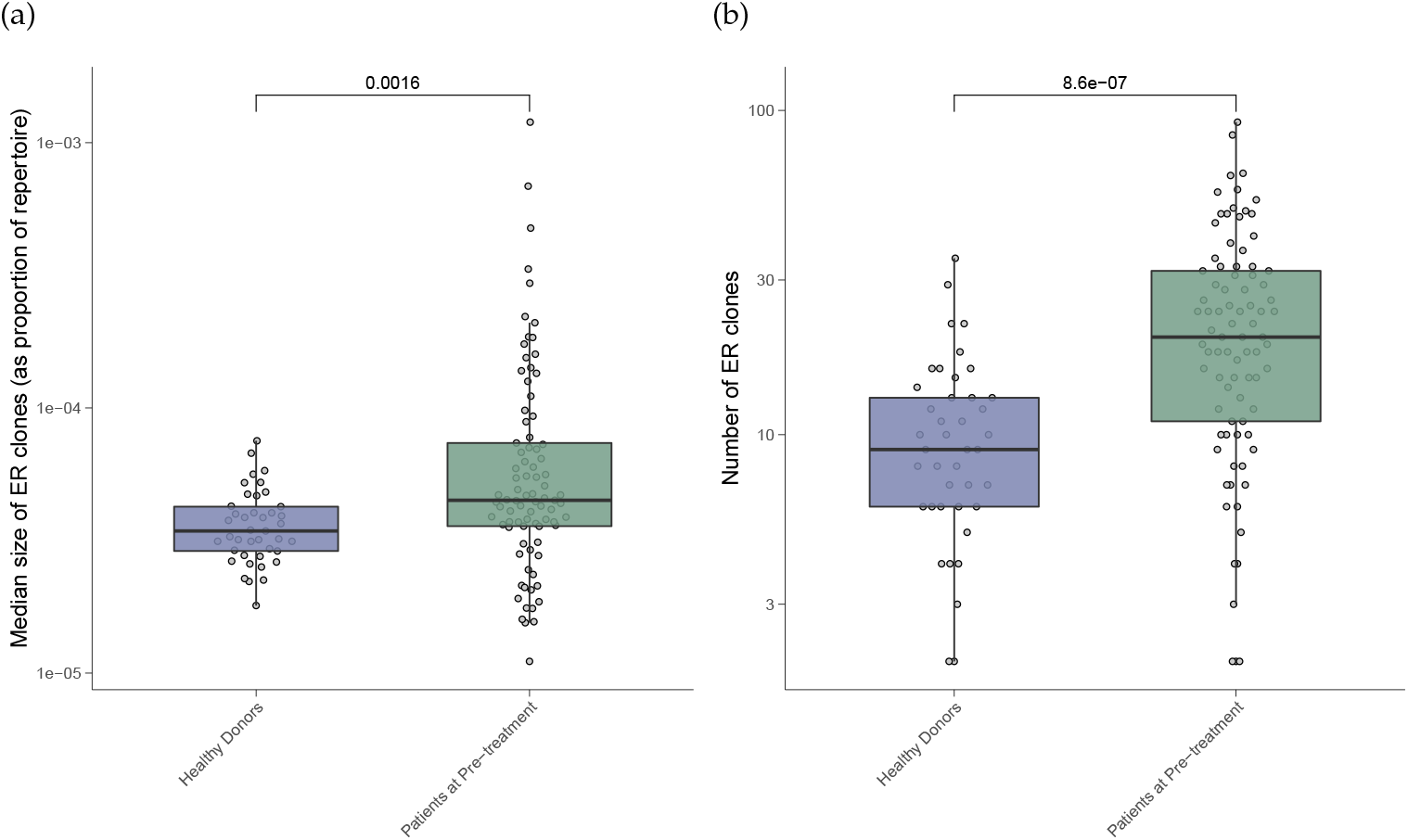
(a) Median size of ER clones in healthy donors and patients pre-treatment. (b) Number of ER clones found in the repertoires of healthy donors and patients pre-treatment. Statistics with two-sided Wilcoxon rank sum test.

**Figure S5.**
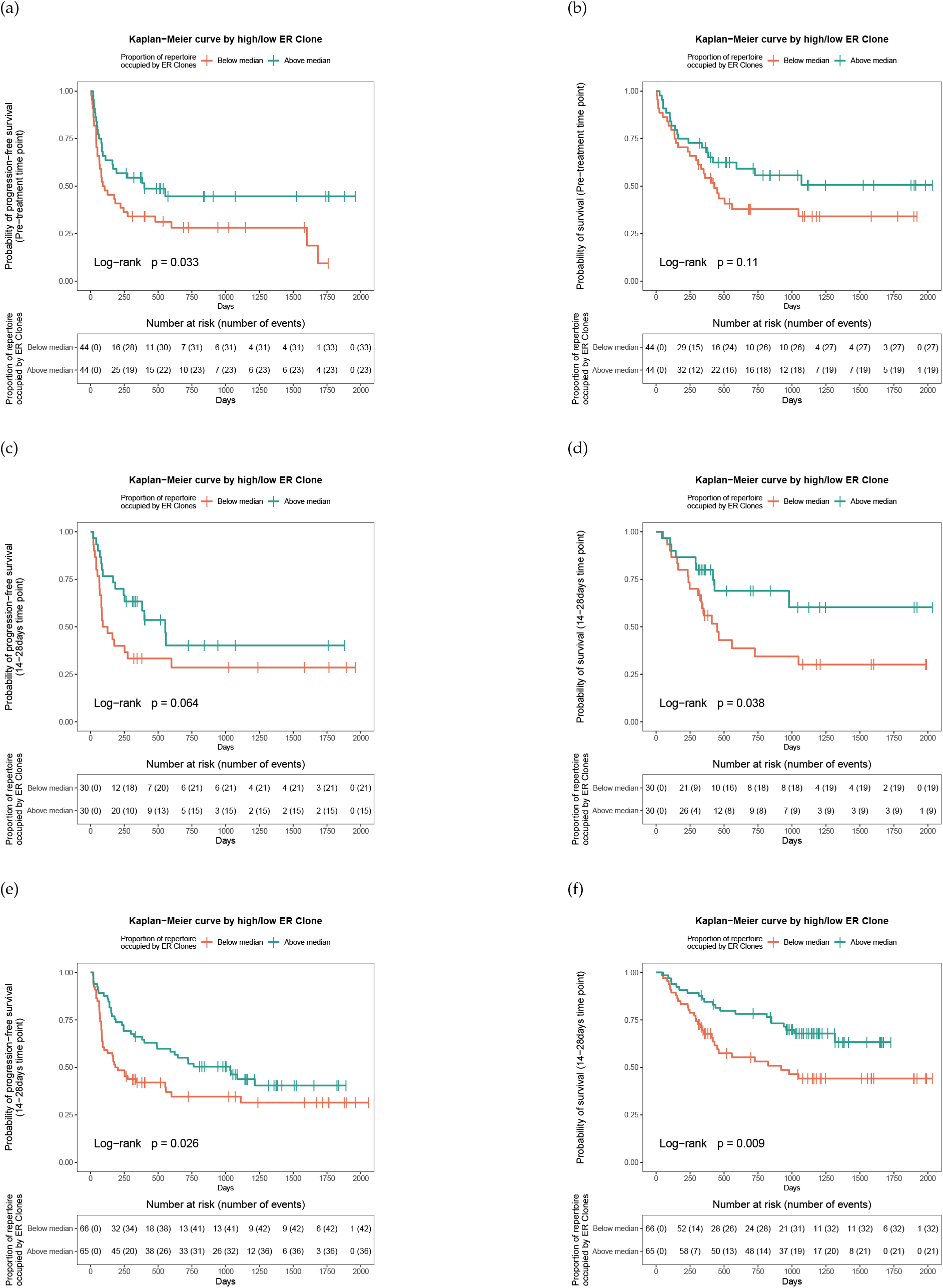
(a) Progression-free survival curves based on above or below median proportion of the repertoire occupied by ER clones at pre-treatment in validation cohort (Fairfax MiXCR). (b) as per (a) but overall survival. (c) Progression-free survival curves based on above or below median proportion of the repertoire occupied by ER clones at 14-28 days post-treatment in validation cohort. (d) as per (c) but overall survival. (e) Progression-free survival curves based on above or below median proportion of the repertoire occupied by ER clones at 14-28 days in pooled discovery and validation cohort (bulk RNAseq data). (f) as per (e) but overall survival. Statistics via Log Rank test.

